# Theoretical Models of Obligate Mutualism to Link Micro- with Macro-Coevolutionary Dynamics

**DOI:** 10.1101/2024.05.16.594543

**Authors:** Carrie Diaz Eaton, Christopher M. Moore

## Abstract

There is a need to link micro- and macro-coevolution to bridge mechanistic theory and observations of micro-coevolutionary change with observations of macro-coevolutionary patterns. This need is particularly conspicuous in theoretical models of obligate mutualism, where phylogenetic matching is the predicted outcome. However, these theoretical models of obligate mutualism create a mismatch with empirical studies of obligate mutualism, which experience extensive phylogenetic discordance. Although environmental variation on geographic scales is often invoked, there are other, non-mutually exclusive mechanisms that can possibly explain genetic diversity and co-phylogenetic patterns in mutualistic communities. In this study, we use a genetic-explicit mathematical model of obligate mutualism that explain host-switching outcomes and, consequently, discordance in cophylogenies. We then explore the role of temporal variation in maintenance of genetic variation (*i.e*., phenology), which can further account for phylogenetic discordance. These insights are possible due to the focus on initial conditions and short-term behavior of model results. This work ultimately supports the continued importance of theoretical work which expands its analysis of outcomes beyond asymptotic behavior.

## Introduction

Coevolution is reciprocal evolution in which interacting species affect each others’ evolutionary trajectories (Thompson, 1982). A theoretical coevolutionary framework positions the magnitude(s) and direction(s) of fitness effects due to species interactions as key in shaping the evolutionary trajectory of each species. Species interactions are typically classified into three categories according to the direction of the non-zero fitness effect of each species—competition (–,–), antagonism (+,–), mutualism (+,+)—that lead to different coevolutionary dynamics. These interactions affect both micro- and macro-evolution of each species, which we will respectively refer to here as micro- and macro-*co*evolution to be explicit about the joint evolutionary dynamics. Micro-coevolutionary theory focuses on understanding genotypic and phenotypic evolution within species. Macro-coevolutionary theory typically focuses on *co*speciation and *co*phylogenesis, the formation and differentiation of clades of interacting species. In this study, we ultimately focus on the micro- and macro-coevolution of the least investigated of these species interactions, mutualism.

Mutualism is a direct, interspecific interaction in which both species’ fitnesses are positively impacted by the interaction. Examples of common mutualisms include plant-pollinator, plant-seed disperser, anthozoans (coral, anemones) and dinoflagellate endosymbionts, and ants and the plants they defend. Ecologically, mutualism is associated with increased growth rates or equilibrium densities at the population level (Vandermeer and Boucher, 1978; Wolin and Lawlor, 1984) and increased diversity and resilience at the community level (Bascompte and Jordano, 2007; Rohr et al., 2014). Theoretical and mathematical models of micro-coevolution of mutualism have explained increased genetic diversity (Stoy et al., 2020) and phenotypic convergence between species and guilds (sets of species performing the same function, such as pollinators) pairs (Thompson, 1989). Models of macro-coevolution have demonstrated how mutualism can both promote and impede diversification and display a wide-range of phylogenetic-tracking (Chomicki et al., 2020, 2019).

Micro-coevolutionary processes serve as mechanistic bases for macro-coevolutionary patterns, and linking the two is important to make inferences between the two scales of evolution (Blasco-Costa et al., 2021; Dismukes et al., 2022). In this study, we consider the simplest case of mutualism—mutually obligate, mutually specialized mutualism; i.e., obligate specialists. Obligate, in contrast to facultative, refers to the case where both sides of the interaction (species, guilds) have a non-positive fitness in the absence of a mutualist; and specialized, in contrast to generalized, refers to the case where one side of the mutualism is composed of one or few species (Chomicki et al., 2020). Obligate specialism occurs in many well-studied systems including yuccas and yucca moths (Pellmyr and Huth, 1994), ants and nest fungi (Schultz and Brady, 2008), figs and fig wasps (Ramírez, 1970), and termites and fungi (Aanen et al., 2009). These specialist interactions are useful for linking micro-processes and macro-coevolutionary patterns because of the simplicity of the one-to-one species ratios. This simplicity allow us to more easily track and make inferences about genotypic and phenotypic patterns at the micro-scale and cophylogenesis at the macro-scale.

Our *a priori* expectation of obligate specialists is they cospeciate, which results in the long term as cophylogenetic concordance between lineages Machado et al. (2005); Page (2003). Prior quantitative trait loci models suggest that trait matching is the only coevolutionary outcome (Kiester et al., 1984; Medeiros et al., 2018). However, empirical observations, even in these relatively simple systems, often show degrees of discordance through patterns of host-switching and host-range expansion (i.e., having positive fitness with new host species while also maintaining its association with the ancestral host) resulting in 1-1 violations (Cruaud et al., 2012; Satler et al., 2019). In fig-fig wasp studies, the existence of hybrid complexes and a “messy” cophylogenetic relationship suggests that the 1-1 “rule’ is infrequently obeyed (Machado et al., 2005; Marussich and Machado, 2007; Yu et al., 2021). Explaining cophylogenetic discordance in these diverse lineages has, consequently, been a macro-coevolutionary research priority for many evolutionary biologists.

The seeming incompatibility of trait matching predicted of existing models and the messy hybrid complexes observed empirically also frames a micro-coevolutionary quest for mechanisms which maintain genetic diversity in populations. Classic genetically-explicit population models assume that heterogeneous environments are abiotic and unresponsive to evolution (Levene, 1953). In the Geographic Mosaic of Coevolution, spatial heterogeneity is invoked to explain how populations adapt to local patches, but genetic diversity in the metapopulation is maintained through gene flow between different patches (Thompson, 2005). Building on this approach, patches are subject to variance due to external forcing (Frank and Slatkin, 1990) or variation in gene flow (Gomulkiewicz et al., 2000). These models also focus on long-term outcomes and equilibrium dynamics.

In this study, we develop biologically-informed mathematical models to investigate the micro-coevolutionary basis to explain both cophylogenetic discordance as well as maintenance of genetic diversity. We first build a mechanistic, genetically-explicit coevolutionary model, in which patches are plant populations that are dynamically evolving in response to pollinator interactions. We then re-interpret our results with a macro-coevolutionary lens. We explore subtle, but key outcome differences when considering a possible genetic basis, which are only observed when considering short-term dynamics. With this model, we can demonstrate a possible genetic basis by which one can observe both fidelity of mutualism that may maintain lineage tracking, as well as host-switching that may lead to more complex macro-coevolutionary patterns.

In the second model, we consider the effect of temporal, rather than spatial, variation in the maintenance of genetic diversity in mutualistic systems. This allows us to explore additional mechanisms that may be involved in maintenance of genetic diversity. More specifically, we consider how phenology can provide a mechanism for genetic diversity to be maintained long-term in pollinator populations.

### Incorporating an Evolving Resource

In the following model we consider expanding a two habitat model with diallelic loci which control preference and adaptation for each niche. In the standard Levine model, populations will maintain diversity driven by the relative habitat availability (Levene, 1953). We consider that the habitat itself responds to the frequency of visitation. This coevolutionary approach reflects the ongoing relationship between plant and pollinator mutualists. For example, plant genetics may determine a chemical profile, which both attracts a certain type of insect and provides a suitable chemical environment for insect oviposition Gibernau et al. (1997); Schiestl et al. (2021). The visitors attracted are also responsible for pollination, directly determining seeds set for the next generation.

As a case study, consider the life cycle of the *Greya* moth and its host plant beginning at the larval stage. The larvae drop to the ground and overwinter in the soil, the adults emerging in the spring. Adult moths find mates on the host plant and copulate, then females deposit their eggs in flowers (Gomulkiewicz et al., 2003). Thereafter, we will refer to this model as H-GM to reference the host-*Greya* moth dynamics.

### Dynamics of Insects

Let *x*_*i*_ be the frequencies of insect genotypes in the pool of adults emerging from soil. As we initially develop this general model, we allow for an unspecified number of alleles and loci, so we use the subscript *i* to refer to insect genotype 1, insect genotype 2, etc. We present a more specific case study with assigned genotypes in the next section. Similarly, let *y*_*m*_ be the frequencies of the plant genotypes at that time. *π*_*im*_ is the preference of an insect with genotype

*i* for a plant of type *m*, and is a complete preference when *π*_*im*_ = 1 and is no preference when *π*_*im*_ = 0. We will use indices *i, j, k* for insects and *l, m, n* for plants.

The frequency of adult insects of genotype *i* found on plants of type *m* is

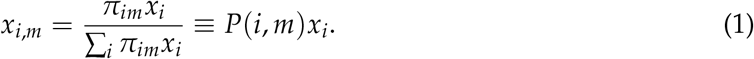

The proportion of the adult insects found on plants of type *m* is

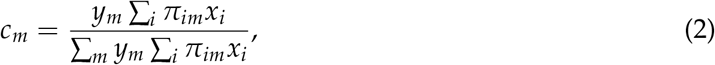

the frequency of insects joining a particular mating pool *m*, normalized and weighted by plant frequency.

We assume random mating on host, with offspring, where *R*(*i, j* → *k*) gives the corresponding offspring frequencies for a given set of parental genotypes. Insects re-assort to lay eggs, and then larvae undergo selection based on their genotype. Therefore the next generation insect frequencies can be calculated by:

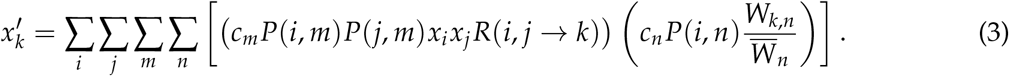

There is no direct competition between larval types. Rather, selection operates weighted to the relative fitness, which could be argued as a form of indirect selection. We can make additional modifications to this model, such as assuming that larvae frequency also affects plant survival or that insects oviposit on the same plant on which they mate. Results for these models are in Appendix A, but the qualitative dynamics are similar.

### Dynamics of Plants

Now consider the genotype frequency change in plants. The probability that a female of type *i* goes to a plant of type *m* for mating and then to a plant of type *n* to lay the eggs (and pollinate the plant *m* with plant type *n*) is *Q*(*i, m*)*y*_*m*_*Q*(*i, n*)*y*_*n*_, where

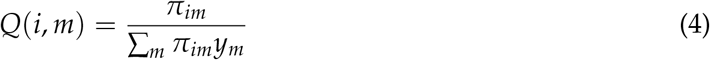

is the probability that an insect *i* visits a particular plant of type *m*.

Thus, the frequency of plant *l* produced as a result of mating of plants *m* and *n* is

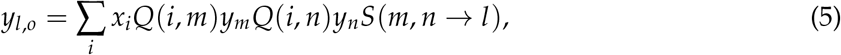

where *S*(*m, n* → *l*) is the corresponding segregation probability.

To account for the fact that the host plants are perennial we assume that only a proportion *β* of plants is replaced each generation by the offspring. Therefore *β* may also be interpreted as 1 over the average number of years in a plant lifespan. Then

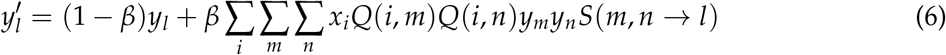

1 − *β* is the proportion of plants that are perennial from the last generation of plants. This may also be interpreted as the proportion that were randomly wind pollinated (and thus have no change in frequency).

Code to reproduce all simulations and figures for this and the following models can be found here: https://github.com/mathprofcarrie/mutualism.

### Diallelic Haploid Host-*Greya* Moth Model

We assume for insects, there are two loci which control adaptation (locus A) and preference (locus B). These diallelic haploid loci can produce four genotypes (1:**AB**, 2:**Ab**, 3:**aB**, and 4:**ab**). For plants we assume a single diallelic locus with two genotypes 1:**C** and 2:**c**. The recombination rate between the loci is *r*.

In this scenario, **A** larvae are optimally adapted to **C** plants and **a** larvae are optimally adapted to **c** plants. Larvae born on the plant they are optimally adapted to have a fitness of 1, otherwise they suffer a selective effect of 1 − *s*. The selection coefficient, *s*, satisfies 0 ≤ *s* ≤ 1. Similarly, **B** insects preferentially visit **C** plants and **b** insects preferentially visit **c** plant. The probability of visitation to a preferential plant is *π* = (1 + *ϵ*)/2, whereas the probability of visitation to a “mismatched” plant has a smaller probability, *π* = (1 − *ϵ*)/2. The parameter *ϵ* can be interpreted as the bias of an insect towards a particular plant choice, and 0 ≤ *ϵ* ≤ 1. When *ϵ* = 0, the insect has no preference. When *ϵ* = 1, the insect will choose the preferred plant every time, never making a “mistake.” See Table 1 for a list of loci, variables and parameters under consideration in the this model.

**Table 1:** List of H-GM model loci, variables, and parameters.

| Loci under consideration |  |  |
| --- | --- | --- |
| Locus **A** | Adaptation in insect for plant |  |
| Locus **B** | Preference in insect for plant |  |
| Locus **C** | Chemical profile of plant |  |
| Allele Frequency Variables |  | Range |
| $p_1, q_1$ | Frequency of insect allele **A**, **a** | $[0, 1]$ |
| $p_2, q_2$ | Frequency of insect allele **B**, **b** | $[0, 1]$ |
| $y_1, y_2$ | Frequency of plant allele **C**, **c** | $[0, 1]$ |
| Genotype Frequency Variables |  | Range |
| $x_1$ | Frequency of insect **AB** | $[0, 1]$ |
| $x_2$ | Frequency of insect **Ab** | $[0, 1]$ |
| $x_3$ | Frequency of insect **aB** | $[0, 1]$ |
| $x_4$ | Frequency of insect **ab** | $[0, 1]$ |
| $y_1$ | Frequency of plant **C** | $[0, 1]$ |
| $y_2$ | Frequency of plant **c** | $[0, 1]$ |
| Parameters |  | Range |
| $s$ | Selection coefficient | $[0, 1]$ |
| $\epsilon$ | Preference bias | $[0, 1]$ |
| $r$ | Recombination probability | $[0, 1/2]$ |
| $\beta$ | $1/(\text{Avg lifespan of plant})$ | $[0, 1]$ |

### Equilibrium Results

The resulting mathematical system is robustly bistable, as illustrated through the analytic and numerical simulations. This means there are two steady states to which the system could evolve depending on the initial conditions. More specifically, the system will always go to fixation for either the **A, B, C** alleles or the **a, b, c** alleles. This is illustrated in Figure 1 in which the trajectories of 100 numerical runs are plotted for the same set of parameter values, but with random starting initial conditions. Other equilibrium states can only occur when parameters are at their extreme values - for example, when the preference parameter *ϵ* is 0, which leads to random oviposition despite adaptive value. See Appendix A for an analysis of these edge cases.

**Figure 1.**
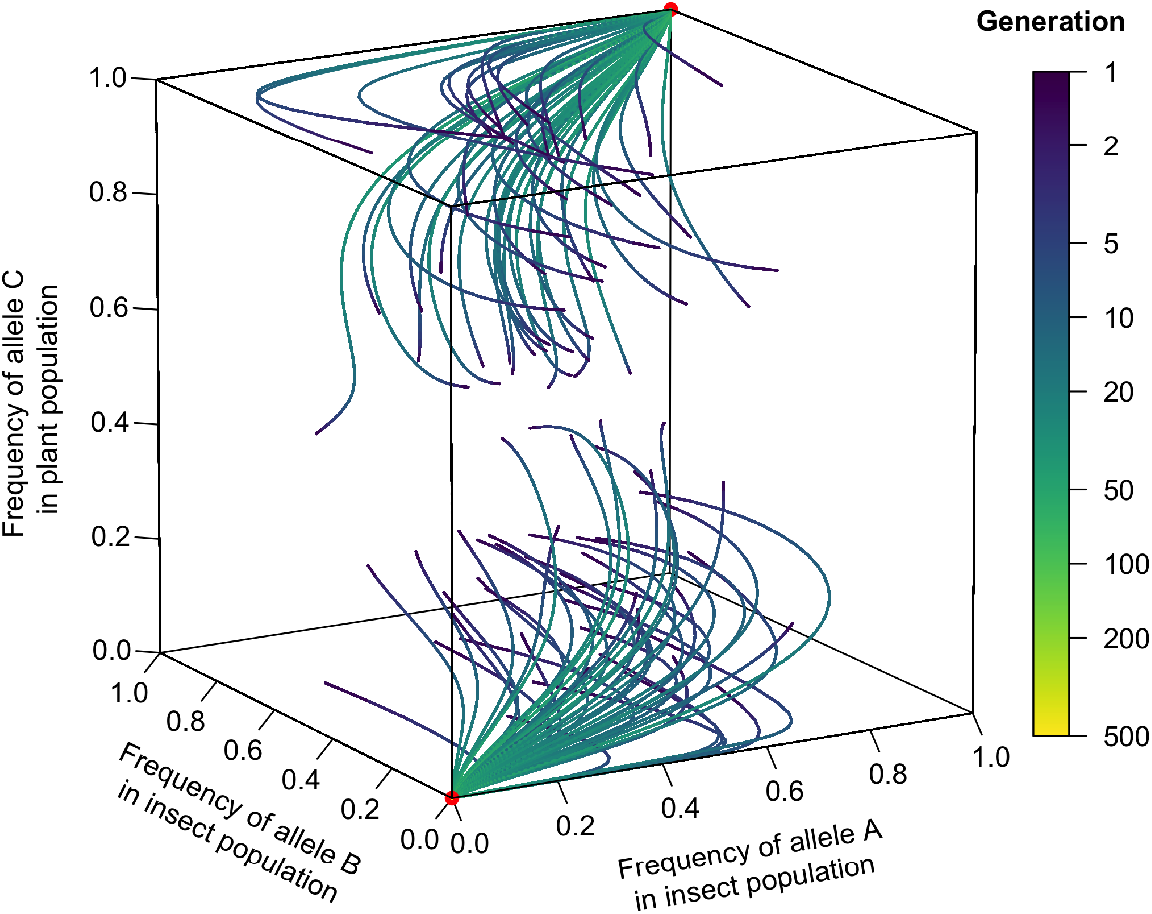
Evolutionary bistability in the host-*Greya* moth model. This phase portrait shows 100 trajectories with 100 random initial conditions under parameter values *ϵ* = 0.5, *r* = 0.2, *s* = 0.6, and *β* = 0.2. The color of the trajectory is gradated with respect to log-scaled time, where the darkest blue is at the initial condition and yellow is at the end of 500 generations. Red dots indicate stable equilibrium at fixation.

### Host-Switching and Short-Term Dynamics

For this model, the short-term model behavior is what connects the micro- and macro-evolutionary processes. For example, in Figure 2, the insect population, despite being initially nearly fixed for the **A** and **B** alleles, evolves to fixation of **a** and **b** alleles. This results from the large initial proportion of **c** plants in the system. Therefore, this system has experienced a host-switch, where the insects evolve to adapt to the more plentiful resource type. Because **c** plants are the most available, the insects that prefer the **c** plants (**Ab** and **ab**) are going to have larger frequencies in the next generation due to mating in higher frequencies. **Ab** will have a selective disadvantage on **c** plants, so the insect genotype that eventually prevails is **ab**.

**Figure 2.**
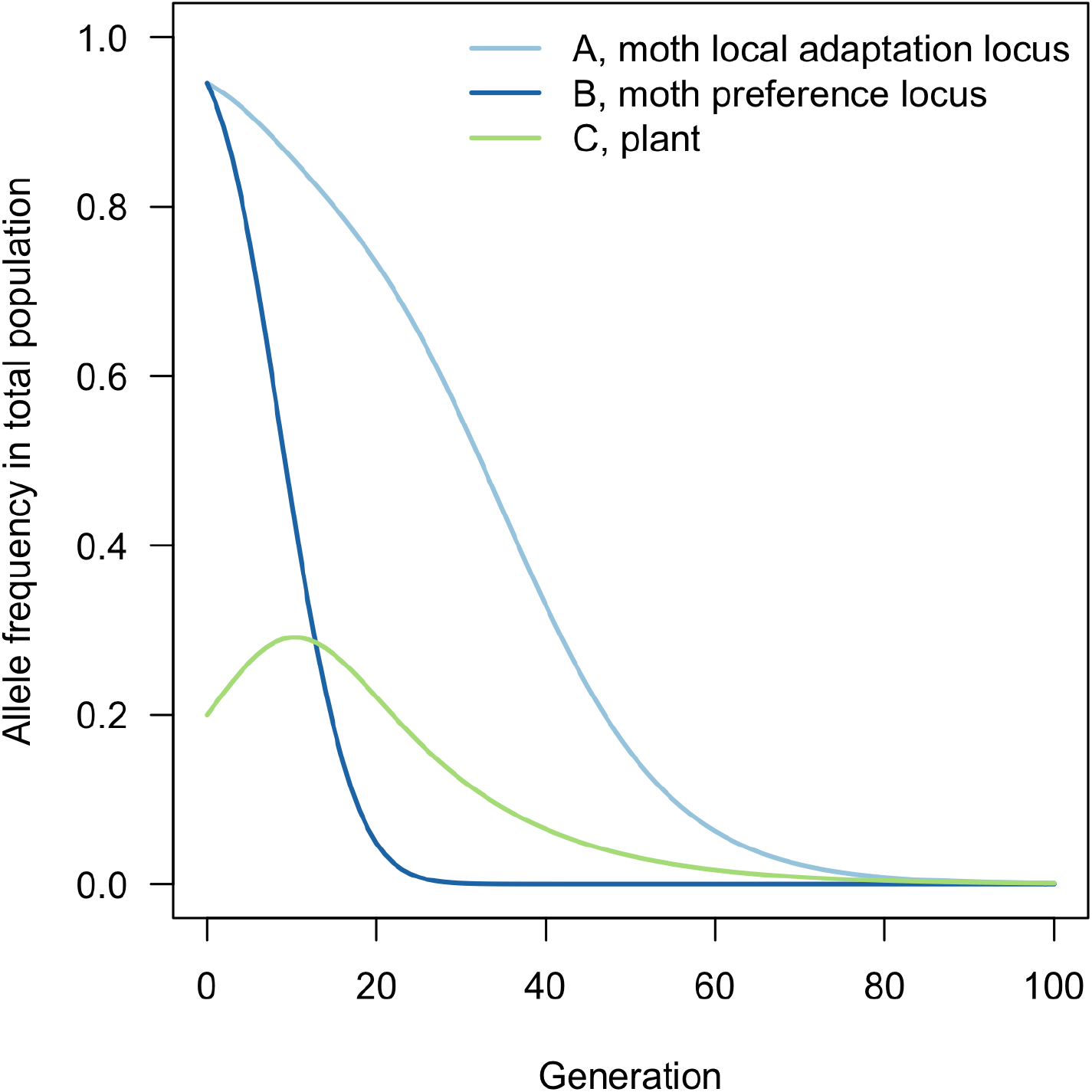
In this simulation, the population initially carries more alleles for adaptation and preference for the **C** plant. However, the most common plant is type **c**. The outcome is for the insect population to rapidly shift in allele frequencies towards exclusively **a** and **b**. In this scenario, the high relative frequency of an alternate plant type drives host-switching in the population. Parameter values *ϵ* = 0.2, *r* = 0.1, *s* = 0.1, and *β* = 0.2.

To gain more intuition into how prevalent host-switching is and how initial conditions affect long term behavior, we look to analytical results. We use a weak selection approximation which assumes that *ϵ, s* ≪ *r* so that higher order terms are negligible Gavrilets (2004). The following equations result, where *q*_1_ = 1 − *p*_1_, *q*_2_ = 1 − *p*_2_, and *y*_2_ = 1 − *y*_1_.

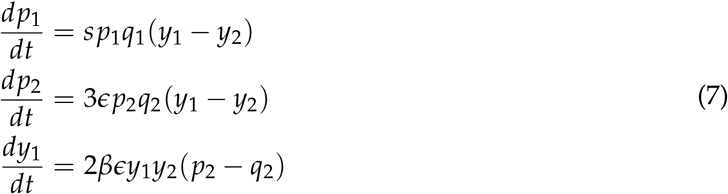

This model reduction can be used to derive the equation of the line which divides the basin of attraction for each fixed equilibrium (the purple line in Figure 3). From there, we can quantify the genotypic space that would result in a host-switch to a new plant type (in this case, about 26%) versus that which is robust to invasion and maintains its dominant association.

**Figure 3.**
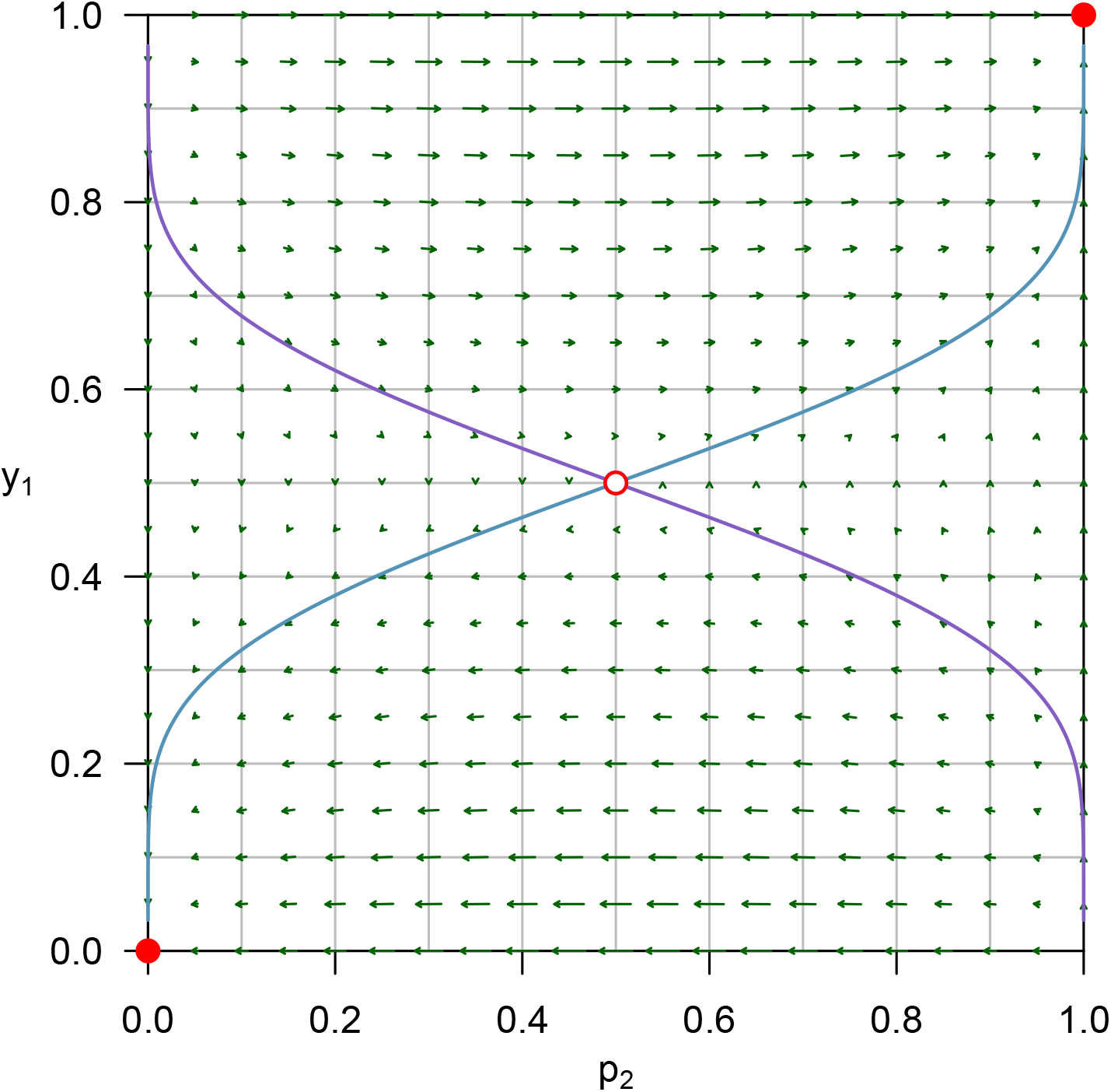
Phase portrait of *p*_2_ vs *y*_1_ for Equation 7 for *ϵ* = 0.1 and *β* = 0.2. Arrows indicate where a trajectory with initial conditions at that point will go. Red dots indicate stable fixed points and the unfilled red circle indicates an unstable equilibrium at (.5,.5). Purple and blue lines indicate the nullclines which delineate the basin of attraction.

### Discussion of the Host-Greya Moth Model

Analytic results show the cases of fixation of **AB** and **C** genotypes or **ab** and **c** genotypes are stable equilibria to which the system evolves long term for realistic parameter conditions. A few exceptions exist as a result of special cases discussed above. Interestingly, numerical simulations and analytical results from model approximations show host-switching as a possible outcome—where a system that has a majority of one type of insect evolves to exploit a dominant plant population of the other type. This agrees with biological observations about this system (Gomulkiewicz et al., 2003; Thompson, 2005) in which host-switching can occur rather easily. Whether or not a host-switch occurs is influenced mostly by the parameter *β* which controls how fast the plant population responds to matching insect abundance. It is less affected by the relative preference insects have for their matching plant, but this observation is not captured in the reduced model.

A complete lack of preference leads to a long-term maintenance of genetic variation in all alleles in the population. This may seem counterintuitive, because it has been conjectured that cases of coexistence in these mutualistic systems are a result of strict preferences (Kerdelhuè, 1997). However, if insects instead oviposit randomly in plants, then the plant frequencies remain unchanged. This essentially converts the system to a two resource problem with only the insect evolving, and thus maintains genetic variability in the population.

Maintenance of genetic variation in this *Greya* moth system is more likely due to a Geographic Mosaic of Coevolution (Thompson, 2005) rather than a complete lack of pollinator preference. However, in many plant-pollinator systems, insects do not show significant correlation between adaptation and preference (Agosta, 2006), and strict preferences can both facilitate and prevent speciation (Gavrilets, 2004). This lack of preference for host may contribute to observed instances of more than one species of wasp ovipositing on the same fig as well as explaining the large amount of hybridization among fig types. We next turn our attention to such systems as we focus on factors that may promote long-term, but not infinite maintenance of diversity.

### Temporal Dynamics

The only micro-coevolutionary pattern not observed in the prior example is duplication, which, from a macro-evolutionary perspective, would maintain a mutualism with a more than one species partner at the same time. In fact our model evolves away from maintenance of such genetic diversity and rapidly towards fixation. This makes the observation of non 1-1 relationships less likely, consistent with the *Greya* inspirational model system. The most common explanation is to zoom out to consider the the [meta-]population and its variations over a broader geographic scale Thompson (2005). We suggest an additional possibility inspired by the reproductive nature of some mutualistic plants—temporal asynchrony due to phenology.

In *Ficus-Courtella wardi* (Wade, 2007), the male phase of the fig from which females emerge, laden with eggs, must overlap with the female phase (the start of a new floral generation of another fig) so that the fig wasp can pollinate the fig and oviposit her eggs. These overlapping generations and the significant delay from seed to flowering maturity could delay plant evolution towards pollinator preference. Thereafter, we will refer to this model as F-FW to reference the inclusion of temporal dynamics inspired by the life histories of some well-studied fig-fig wasp systems.

### Pollinator Dynamics

Consider a small pollination network of only two populations of figs that flower asynchronously from each other—when one population is in male phase, the other is in female phase. This implies that fig wasps have only two generations in the time that it takes for both fig populations to go through their male and female phases; see Figure 4.

**Figure 4.**
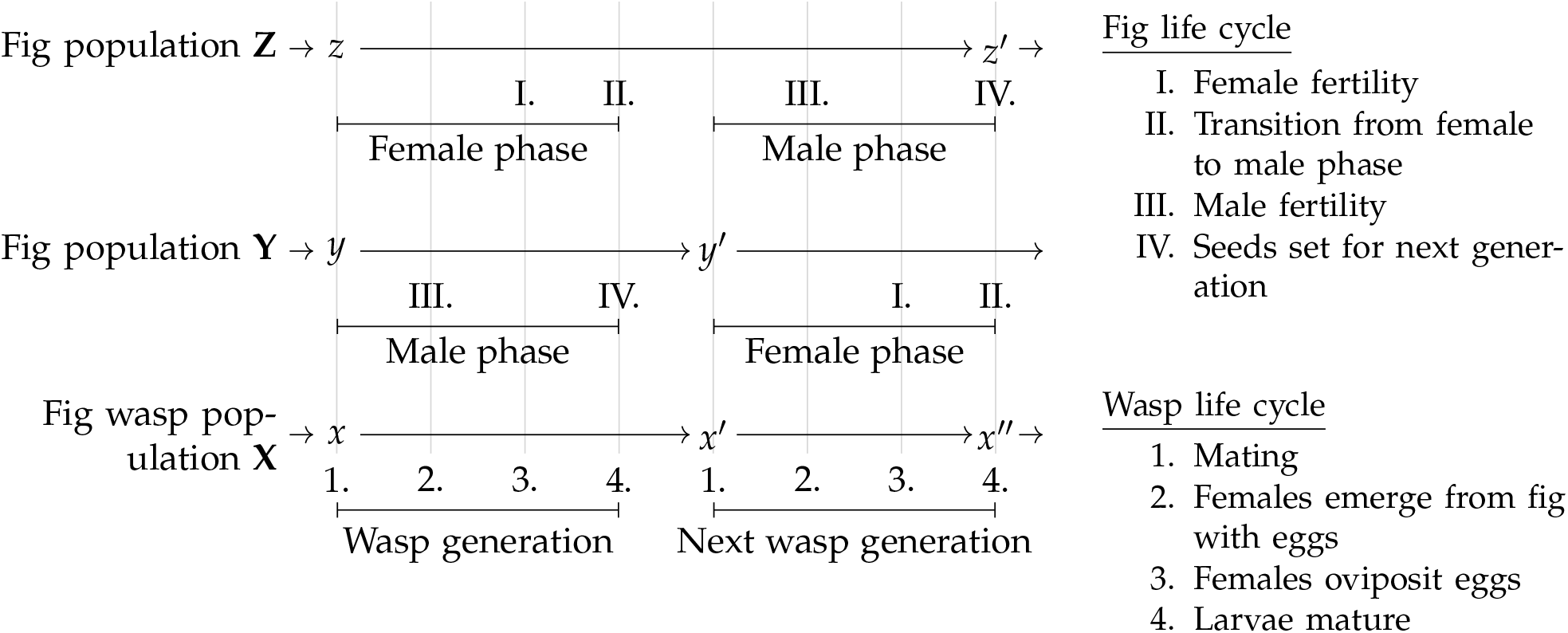
Model life cycle of insect generations and the overlapping and asynchronously flowering plant populations **Y** and **Z**.

We can use prior methods to describe genotype frequencies during all parts of the fig and fig wasp life cycle. While a specific diallelic haploid model is considered for analysis in the next section, we begin building the general model first, with the more complicated life history stages incorporated (see Figure 4 as an example shapshot of stages in the F-FW model). Wasps mate (population **X**, stage 1), adult wasp females emerge from a male phase fig in one of the populations (**X**, stage 2; fig population **Y**, stage III), and oviposit into a female fig in the other population (**X**, stage 3; fig population **Z**, stage I). At this point, the expected genotype frequencies among larvae (of type *k*) of insects maturing in the male phase fig (of type *n*) in the **Y** population of plants can be written as:

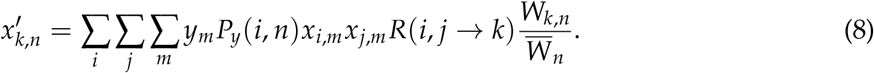

Note that *ϵ* is not explicit in this equation, but plays the same role as in the H-GM model—a measure of choosiness that is embedded into the probability of pollinator visitation function, and the equation above is analogous to Equation 3.

These larvae mature (**X**, stage 4) as the fig transitions to male phase (**Z**, stage II). Note that a full generation life cycle for wasps has completed, and a second generation is required to complete a full flowering cycle for figs. Within the **Z** fig, wasps mate (next generation **X**, stage 1). Then female wasps emerge from the now male phase **Z** population of figs (next generation **X**, stage 2; **Z**, stage III) to oviposit eggs into a plant in the **Y** population of plants which is now in female phase again (next generation **X**, stage 3; next generation **Y**, stage I). Larvae mature in the **Y** population figs (next generation **X**, stage 4), and after selection, the genotype frequencies of insects are:

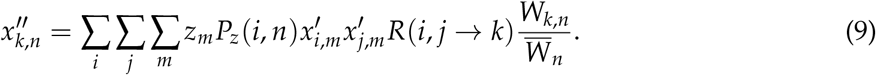

### Host Dynamics

The host species is pollinated when female wasps emerge from a male phase plant in population (**X**, stage 2; **Y**, stage III) and oviposit in a female phase plant (**X**, stage 3; **Z**, stage I). Because this requires a sort of hybridization between populations to be maintained, we assume that flowering time is paternally inherited and assign membership in the Y population to be determined by the flowering time. To account for the fact that figs last more than one generation, we assume only a proportion, *β*, of plants are replaced each generation. The next generation genotypes in population **Y** are then described by the following:

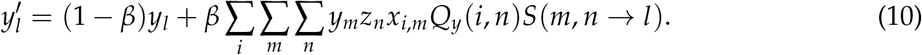

The equations for the next generation of the **Z** population are analogous, but reproduction of **Z** plants (**Z**, stage III; **Y**, stage I) occurs during the second insect life cycle (next generation **X**, stages 2 and 3).

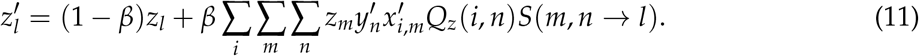

## Considering the Diallelic Haploid Model

In the section below, we analyze the case the of diallelic, haploid F-FW model. This model uses the same variables and parameters of the diallelic haploid H-GM model (see Table 1), except that there are two plant populations, **Y** and **Z**. The frequency of allele **C** in population **Y** is still *y*_1_, but the allele textbfC is also found in population **Z**, with frequency *z*_1_.

### Results of the Fig-Fig Wasp Model

This model variation produces all of the behaviors of the previous model: bistability, some robustness to host-switching, and host-switching (Figure 5). However, it differs in an important feature: the time to reaching the stable equilibrium. If the odd generations of plants are different in composition from the even generation plants, then the insect genetics will oscillate between the two until the odd and even generation plants are genetically similar in composition. For values of *ϵ* nearer to 1, corresponding to very specific preference, the population eventually goes to a fixed state. This is achieved only after several hundreds to thousands of generations of oscillating insect populations affected by the plant populations’ plant type frequencies, though the exact timescale varies with *β*.

**Figure 5.**
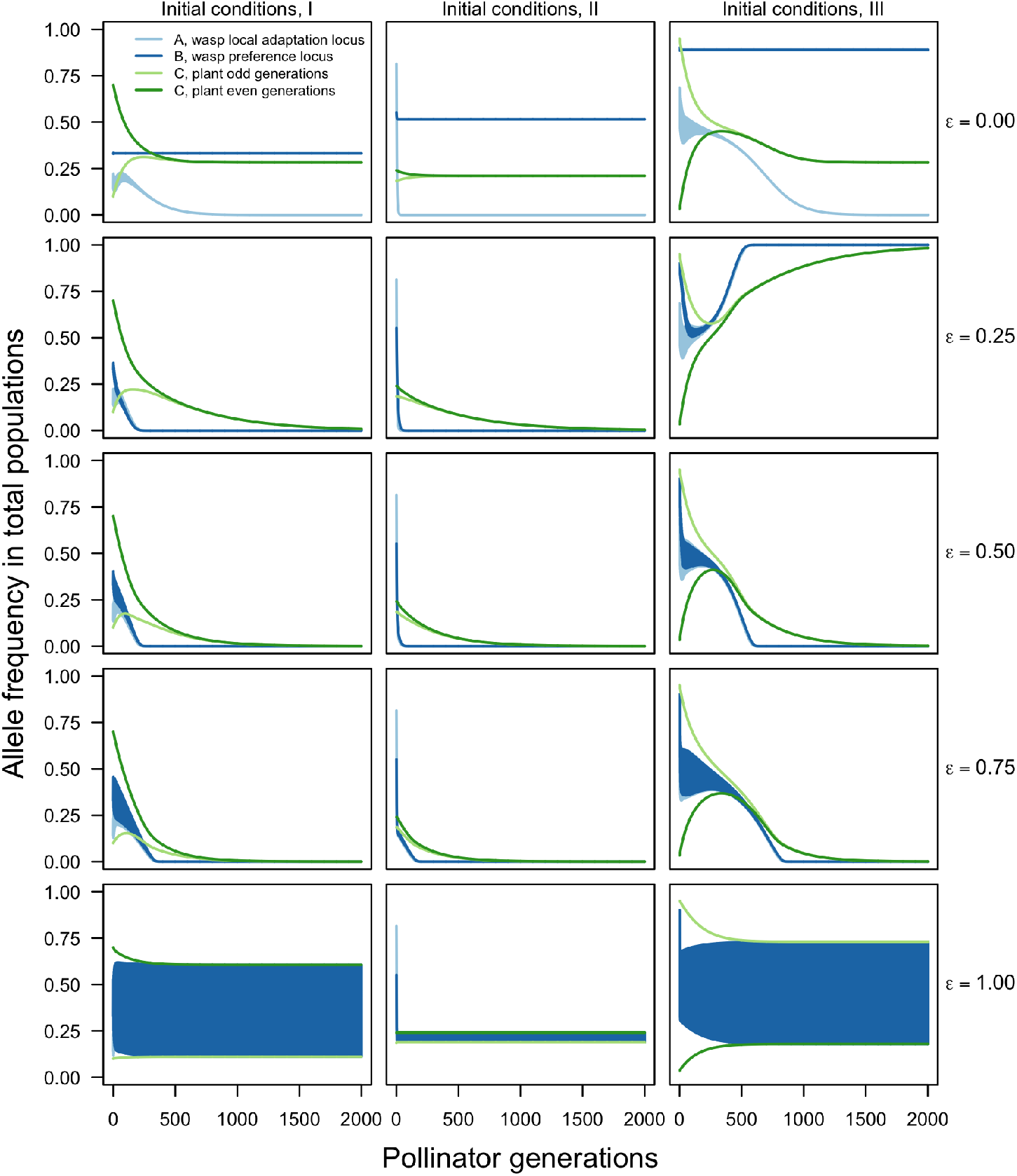
Possible trajectories of the F-FW model over 2000 total pollinator generations. The simulations in each column have the same set of initial conditions, and values for *r* = 0.2, *s* = 0.6, and *β* = 0.02. Each row varies *ϵ* from 0 (top row) to 1 (bottom row) in increments of 0.25. Note that the solid blue areas are actually very rapid oscillations of both the adaptation (light blue) and preference (dark blue) allele frequencies.

### Discussion of the Fig-Fig Wasp Model

Adding temporal variation due to phenology can significantly affect transient behavior. Even under intermediate values of *ϵ* in a simplified model that considers only two plant populations, oscillatory insect and differing plant frequencies between plant populations can persist for thousands of insect generations. A typical fig wasp lineage may rely on 8–10 different fig populations in order to maintain its persistence CM: Citation.. In this case, it is expected that the transient behavior seen in the two-population case will last even longer, perhaps tens of thousands of generations under realistic parameter conditions. This long-term transience may be extremely important in clarifying why so many fig phylogenies are extremely difficult to resolve, pointing to extensive hybridization and confusion when determining historical relationships with pollinators (Weiblen and Bush, 2002).

Looking at larger evolutionary time scales, reproduction of three major coevolutionary phylogenetic patterns in the fig-fig wasp system are observed: robustness of the 1-1 coevolutionary relations between plant and pollinator, host-switching of pollinator to another fig type, and extensive hybridization among figs leading to violations in the 1-1 relationship between fig and fig-wasp. Initial plant type distributions and magnitude of preference bias, *ϵ*, may determine which of these scenarios is observed. A study of fig and fig wasp species pairs in the Ogasawara Islands revealed that *F. nishimurae* and the Higashidaira type selected their own host fig odors significantly more often, while wasps of *F. boninsimae* did not show specific preference for a particular fig odor (Yokoyama, 2003). This preference bias parameter does vary *in situ* and may explain some major variations in evolutionary trajectories in different fig-fig wasp systems.

## Conclusion

It is imperative to link micro- and macro-coevolution to bridge mechanistic theory and observations of micro-coevolutionary change with observations of macro-evolutionary patterns. Because macro-coevolutionary patterns of mutualism tend to be thought of as extreme 1-to-1 or completely diffuse, early theoretical models of mutualism tend to be more teleological and reinforcing the extreme 1-to-1 or diffuse patterns of cophylogeny Kiester et al. (1984). For example, pointing to environmental variation as a driver for plant diversity allows us to maintain 1-to-1-ness at a finer spatial scale (*e*.*g*., (Gomulkiewicz et al., 2003; Michaloud et al., 1996; Rasplus, 1996; Weiblen, 2002)). However, since most cophylogenetic patterns lie somewhere between 1-to-1 and diffuse, we need to use inductive and mechanistic reasoning to explain those patterns and view the edge extremes as rare and specialized cases (Machado et al., 2005; Michaloud et al., 1996).

Together our models hint towards the micro-evolutionary mechanisms that may shape macro-evolutionary trajectories. For example, when pollinator populations resist a potentially newly plant abundant type, this may promote their continued lineage tracking together. However, in our models pollinators or plants do sometimes switch, adopting to a more prevalent partner type. This is well supported also by studies on *Greya* moths and their *Lithophragma* hosts (Thompson, 2005). Finally, our F-FW models illustrated a third pattern—maintenance of genetic diversity for thousands of generations. The temporal asynchrony in phenology which maintains diverse partnerships may be one of the drivers of hybrid complexes observed in the fig-fig wasp mutualistic system (Machado et al., 2005).

Because previous theoretical work on mutualism suggested long-term fixation, the theoretical focus of coevolution moved to antagonistic interactions to explain maintenance of genetic diversity (*e*.*g*. Nuismer et al. (2007); Nuismer and Thompson (2006). However, we find theoretical support for mutualism as sufficient to explain both host switching and maintenance of genetic diversity through long time scales, as illustrated through simulations at various initial conditions and phase plane analysis of model approximations. These important theoretical results are missed if the focus of mathematical model analysis is solely on stable asymptotic dynamics (Hastings, 2001; Hastings et al., 2018).

Recent discussion on the importance of considering transient dynamics has primarily focused on observing different types of stable, cyclic or chaotic states, the overall philosophy which cautions against over-reliance on asymptotic analysis applies to our findings as well (DeAngelis and Waterhouse, 1987; Hastings et al., 2018). More recently, the conversation has focused to specifically define long transients to include change slow relative to other evolutionary forces (Morozov et al., 2020). Models which include spatial or temporal structure, like the F-FW, may lengthen the time that transient behavior is observed (Hastings et al., 2018; Morozov et al., 2020).

## Acknowledgments

Omitted for anon review

## Appendix A Simple Coevolution Model

There are a large number of plants on which mating and oviposition occur, mathematically we assume infinitely many, and plants are pollinated by ovipositing females.

### Dynamics of Insects

To understand how insect genotype frequencies change over time, consider the change that occurs in each generation as a difference equation. Let *x*_*i*_ be the frequencies of insect genotypes in the pool of adults emerging from soil and let *y*_*m*_ be the frequencies of the plant types at that time. *π*_*im*_ is the preference of an insect with genotype *i* for a plant of type *m*. We will use indices *i, j, k* for insects and *l, m, n* for plants.

The frequency of adult insects of type *i* found on plants of type *m* is

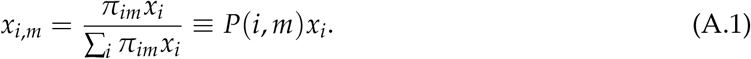

where

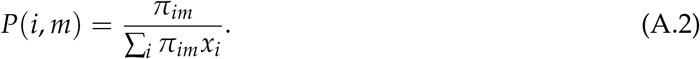

can be interpreted as the preference of insect *i* for plant *m* relative to the average preference of insects for this plant.

The proportion of the adult insects found on plants of type *m* is

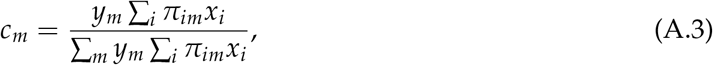

Assuming random mating on host, the frequency of mating pairs formed by females *i* and males *j* on a plant of type *m* is

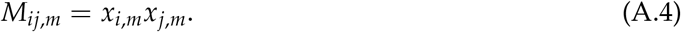

The frequency of eggs with genotype *k* produced by pairs (*i, j*) mating on a plant of type *m* is *M*_*ij,m*_*R*(*i, j* → *k*), where *R* gives the corresponding offspring frequencies for a given set of parental genotypes.

We assume the contribution of each mating pool (*i*.*e*. plant type) to the overall offspring pool is equal to the proportion of insects that came to the pool. Thus, the proportion of eggs with genotype *k* and carried by mothers with genotype *i* in the whole set of eggs before oviposition is

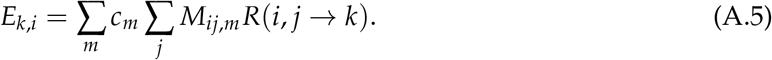

The frequency of eggs with genotype *k* oviposited on a plant of type *n* (by all females *i*) is

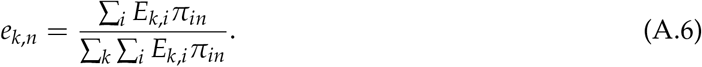

After selection on a plant of type *n*, the frequencies of larvae dropping to the soil from that plant are

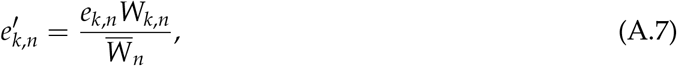

where *W*_*k,n*_ is fitness, *i*.*e*. viability, of genotype *k* on a plant of type *n* and 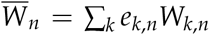 is the average fitness of the population on a plant of type *n*. We assume that the contribution of each larva pool (*i*.*e*. plant) to larvae is equal to the proportion of the eggs deposited on the plant. Note that variations on this assumption will be explored in a later model. The frequencies of the insect genotypes in the pool of adults emerging from soil in the next generation are

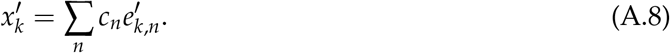

The difference equation for insect genotypes in the next generation is then

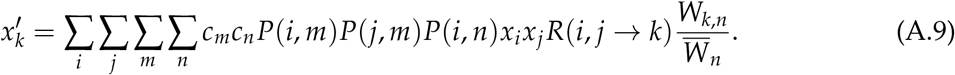

### Dynamics of Plants

Now consider the genotype frequency change in plants. The probability that a female of type *I* goes to a plant of type *m* for mating and then to a plant of type *n* to lay the eggs is

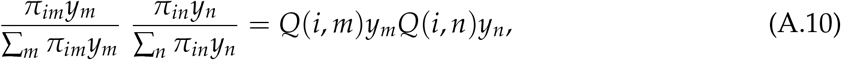

where

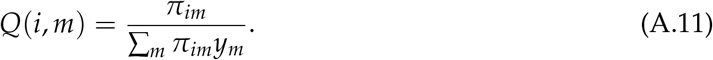

By doing this the female pollinates a plant *n* by pollen from plant *m*. The term *Q*(*i, m*) can be interpreted as the probability that an insect *i* visits a particular plant of type *m*. Thus, the frequency of plant *l* produced as a result of mating of plants *m* and *n* is

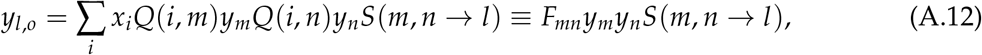

where *S*(*m, n* → *l*) is the corresponding segregation probability (recall there is no recombination since we only consider one plant loci), and the term

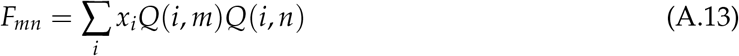

can be interpreted as fertility of mating pair *m* and *n*.

Finally, to account for the fact that the host plants are perennial we assume that only a proportion *β* of plants is replaced each generation by the offspring. Therefore *β* may also be interpreted as 1 over the average number of years in a plant lifespan. Then

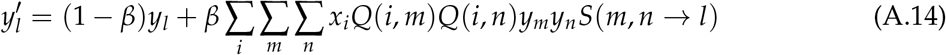

### Plant Resource Dependent Model

In creating our initial model, it is assumed the frequency of adult insect types in the next generation is proportional to the larva frequency types surviving on each plant and the frequency of egg types laid on the each plant. This assumes there is no plant resource limit or larval competition on plants. In a variation of that model, we assume the contributions of larva types remaining after viability selection are weighted in the next generation by the frequencies of the plant they are on. This changes Equation A.8 to

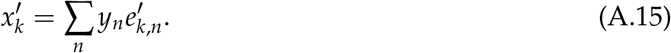

Then the equation for the next generation of adult insect is

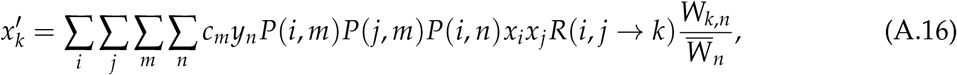

which can be compared to Equation A.9 above:

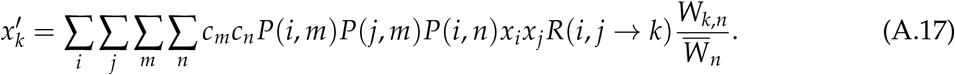

### Model without Re-assortment

To further simplify the above models, one of the multiple moth assortments to its host plant is eliminated. If multiple re-assortment merely shuffles insects but maintains the same frequencies on each plant, then mathematically we could delete one of the trips the female insect makes. In this model variation, instead of re-assorting to lay eggs, she lays eggs on the plant she mates on. In doing so, analytical tractability is improved without changing the qualitative behavior. The modification is made to the H-GM plant resource dependent model. This eliminates the *c*_*m*_ term, because females no longer have to re-assort to lay eggs:

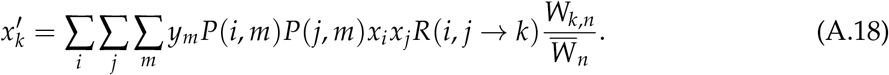

### Analytical Results

#### Assumptions

For the following numerical and analytic analysis, we consider the case where each locus is diallelic. Let *x*_*i*_ (*i* = 1, 2, 3, 4) be the frequencies of four insect genotypes, **AB, Ab, aB** and **ab**, respectively, in the pool of adults emerging from soil. Let *y*_*m*_ (*m* = 1, 2) be the frequencies of the two types of plants (**C** and **c**, respectively) during the insect mating period. A list of insect and plant genotypes matched up with indices is in Table A.1.

**Table A.1:** A list of insect and plant genotypes in the H-GM model listed by index.

| Insect Genotypes |  |
| --- | --- |
| Index | Genotype |
| 1 | AB |
| 2 | Ab |
| 3 | aB |
| 4 | ab |
| Plant Genotypes |  |
| Index | Genotype |
| 1 | C |
| 2 | c |

Recall *π*_*im*_ is the relative preference of an insect with genotype *i* for a plant of type *m*. We consider the case where an insect having allele **A** encountering plant of type **C** will chose that plant with probability *π*_1,1_ = *π*_2,1_ = (1 + *ϵ*)/2, but will chose a plant of type **c** with probability*π*_1,2_ = *π*_2,2_ = (1 − *ϵ*)/2 (vice versa for an insect carrying the **c** allele). *ϵ* can be interpreted as the bias of an insect towards a particular plant choice and 0 ≤ *ϵ* ≤ 1. When *ϵ* = 0, the insect has no preference and chooses whomever it encounters first. When *ϵ* = 1, the insect will choose the matching profile every time and will never make a “mistake.” Likewise, insect larvae born on a matching plant type will have higher fitness.

At the local adaptation loci an insect may have either allele **A** or **a**, which are best adapted to plant types **C** and **c**, respectively. We consider the case where a larvae laid on a matching plant to which it is adapted will have fitness *W* = 1, and a larva developing on a non-matching plant will have fitness *W* = 1 − *s. s* is referred to as the selection coefficient and 0 ≤ *s* ≤ 1.

#### Analytical Methods

The model introduced above is a non-linear discrete dynamical system. Time is described by generations, and each generation’s genotype frequencies are calculated from the previous generation’s. To predict the outcome of the system after many generations of evolution, equilibria and their stability are examined. Stable equilibria are of particular interest, because the system will evolve toward a stable equilibrium in the long term. Full analytical investigation of each model is presented in Appendix A.

Classic 1-1 co-evolution with one’s evolutionary partner, host-switching and speciation or the maintenance of genetic variation are the primary points of interest. For classic co-evolution and host-switching to occur, fixation of each set of matching alleles (**A, B**, and **C** or **a, b**, and **c**) must be a stable equilibria for the system. Whether cospeciation or host-switching is taking place is inferred from initial conditions. Thus both numerical simulations and analytic work are performed to determine where the basin of attraction is for each fixed equilibrium when they are both stable. This means that one can determine how the system behaves long term based on the initial conditions. For the case of speciation, if the polymorphic state is stable, it is possible for both types of plants to co-exist in the system.

#### H-GM model results

Twenty equilibria emerge from the H-GM model, sixteen of which are listed in Table A.2 and four additional biologically unrealistic equilibria, which are not listed. The full analysis proving the stability of the equilibria is presented in the Appendix of Diaz Eaton (2013).

**Table A_2.**
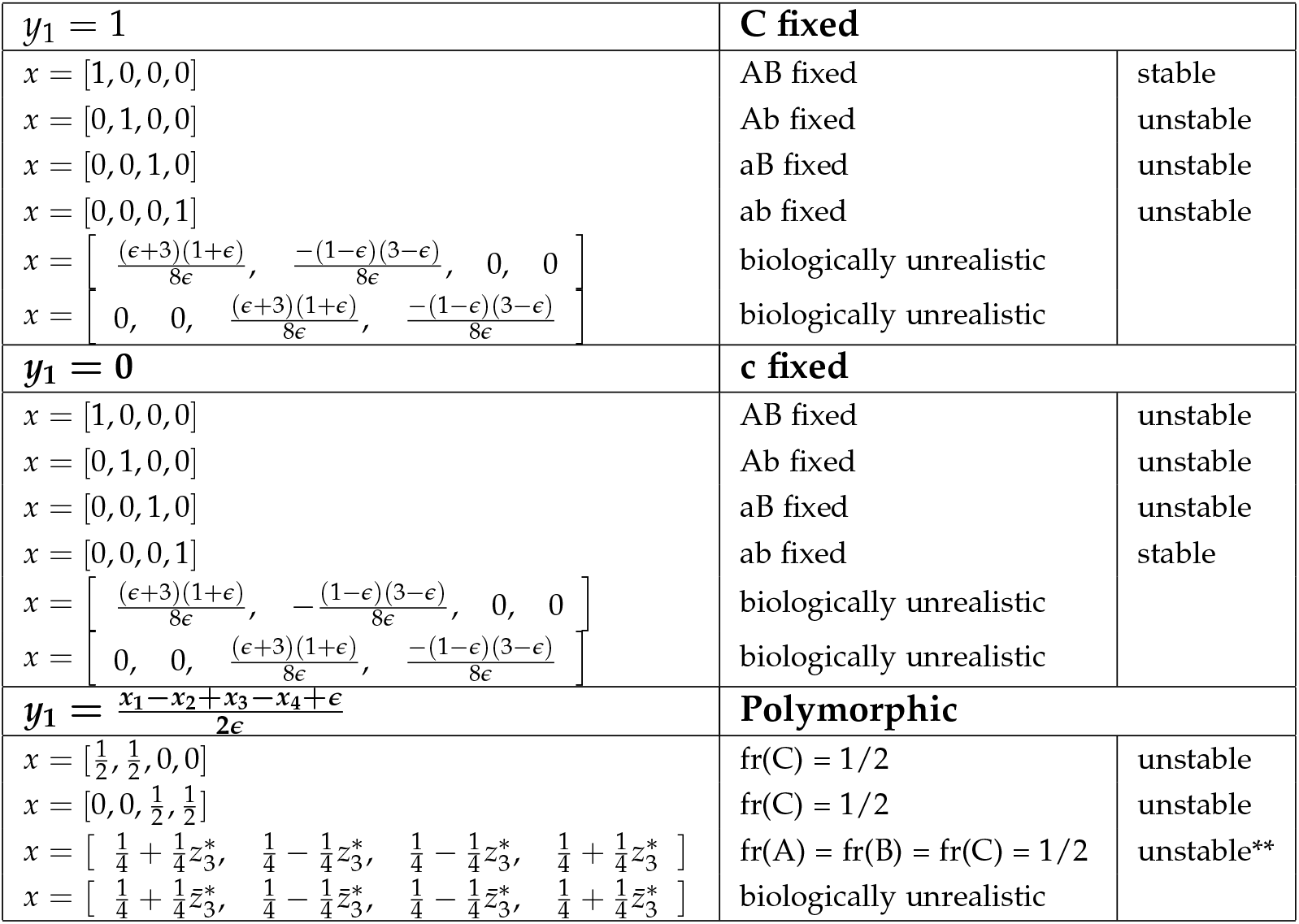
All equilibria are either always stable or always unstable for all biologically realistic parameters unless otherwise noted. **Note, numerical simulations indicate that this point seems to be unstable, however, this result is not shown analytically. 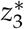 and 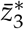, are +/− solutions to a quadratic equation and are functions of *r, s,* and *ϵ*; the exact form is shown in Appendix of Diaz Eaton (2013).

The system is bistable, as illustrated through the analytic and numerical simulations. This means there are two steady states to which the system could evolve depending on the initial conditions. More specifically, the system will always go to fixation for either the **A, B, C** alleles or the **a, b, c** alleles. This is illustrated in Figure 1 in which the trajectories of 100 numerical runs are plotted for the same set of parameter values, but with random starting initial conditions.

#### Approximation of H-GM Model

To gain more intuition into the model and its behavior, a weak selection approximation is used. Here, we assume that *ϵ, s* ≪ *r* so that higher order terms are negligible. Below are the results for approximations of all models under consideration. Note that this assumption will result in *D* = 0, so the transformation that results in variables *p*_1_, *p*_2_, *y*, and *D* will be reduced to only 3 variables. Recall that *D* = 0 was satisfied in the full models’ equilibria, so this simplification will maintain important aspects about the equilibria.

Performing the above approximation, the following equations result, where *q*_1_ = 1 − *p*_1_, *q*_2_ = 1 − *p*_2_, and *y*_2_ = 1 − *y*_1_.

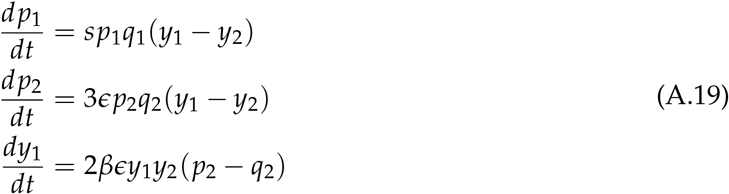

Note: the rate of change in *p*_1_, the frequency of the locus controlling adaptation, is proportional to the selection coefficient, the heterogeneity or variance at the adaptation locus, and the difference in plant type frequency. If there are more of plant type 1 than 2, then the frequency of those adapted to plant 1 will increase. If there are more of plant type 2 than 1, then the frequency of those adapted to plant type 1 will decrease (and so those adapted to plant type 2 will increase). Likewise, the rate of change in *p*_2_, the frequency of the locus controlling plant preference, is proportional to the genetic variance of the plant preference locus and to the difference in plant types. This sets the stage for the preference and locus to go to fixation dependent on which type is the most dominant.

The rate of change in *y*_1_, the locus controlling plant type, is proportional to the variance at that locus and to the difference between the frequency of those having a preference for plant type 1 and those having a preference for plant type 2. Therefore relative plant type abundance is influenced by the relative abundance of pollinators that prefer it.

##### Theorem 1.

*The solutions to the set of equations A*.*19 are*

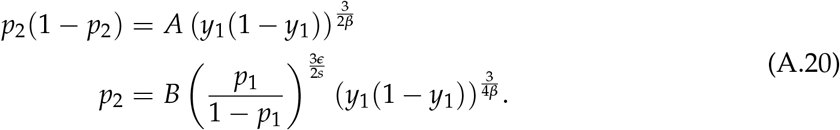

*Proof*. The two differential equations that depend on each other are

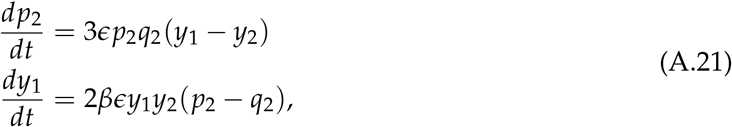

which, when divided, yield

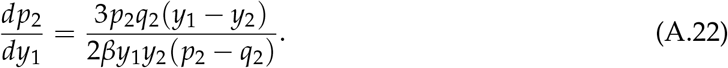

Recall that *q*_2_ = 1 − *p*_2_ and *y*_2_ = 1 − *y*_1_. Then

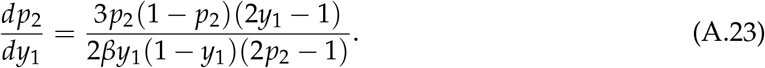

Separate variables and integrate:

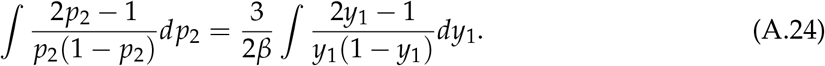

Integration using partial fractions:

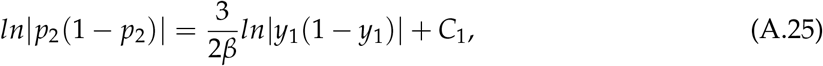

where *C*_1_ is a constant of integration.

This implies that 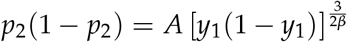, where 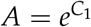.

Now consider

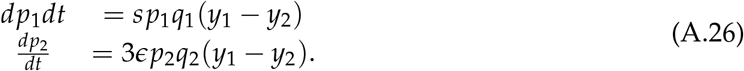

Again, divide and separate by variables:

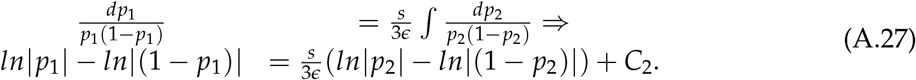

Recall from above 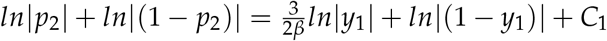,so

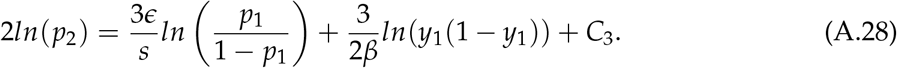

Thus, we conclude

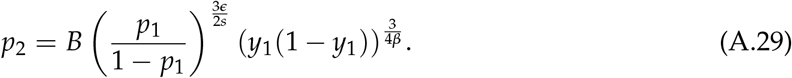

#### Phase Plane

Note that the adaptation locus does not influence the dynamics at the preference locus or the plant type relative abundance. Therefore, analysis of the solutions and phase portrait of *p*_2_ versus *y*_1_ reveals how the initial conditions determine long term dynamics. This phase portrait is illustrated in Figure 3 for a particular parameter set. It shows that the equilibrium point at 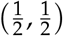 is a saddle point. Also note that the boundary separating the basin of attraction for (0, 0) and (1, 1) is the stable manifold for the saddle point Strogatz (1994).

##### Theorem 2.

*The separatrices, which are the trajectories of the stable manifold of the equilibrium point at* 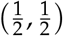 *are*

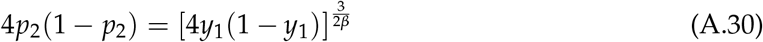

*for* 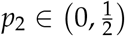 *and* 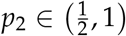

*Proof*. The solutions depicted in the phase portrait in Figure 3 are in the proof of the previous theorem:

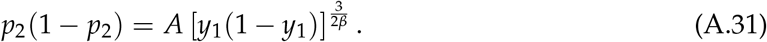

Both trajectories end in *f rac*12, *f rac*12 as *t*− *>* inf. With this condition we get that the basin boundary is:

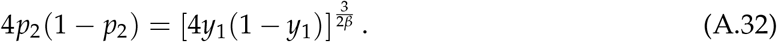

Note that the approximation loses the ability to predict the small changes in the basin boundary due to modifying *ϵ*.

#### Host-Switching

A host-switch to a **C** (type 1) plant is defined as a trajectory with initial condition *p*_2_ *>* .5 and *y*_1_ *<* .5 that goes to the stable equilibrium at (*p*_2_, *y*_1_) = (0, 0). In the example illustrated in Figure 3 where *β* = 0.2, the percentage of area in which this scenario happens is approximately 13.2 percent. This result is attained by integrating the area under the separatrix with the aforemen-dtioned limits. Similarly, since the system is symmetrical, 13.2 percent of the initial condition area will result in a host switch to a **c** (type 2) plant (defining a host-switch to type 2 plant to be trajectory with initial condition *p*_2_ *<* .5 and *y >* .5 that tend towards the stable equilibrium at (*p*_2_, *y*_1_) = (1, 1)).

#### Comparing H-GM Model Variations

##### H-GM Plant Resource Dependent Model

In the full model, a total of twenty equilibria emerge. This model variation is also bistable for the fixed states and the final solution depends on the initial condition. Numerical work confirms the instability of the polymorphic equilibrium whose stability matrix was rather intractable for confirming analytically. Further details can be found in Diaz Eaton (2013).

Using the approximation techniques discussed above for the H-GM plant resource-dependent model, the following equations result:

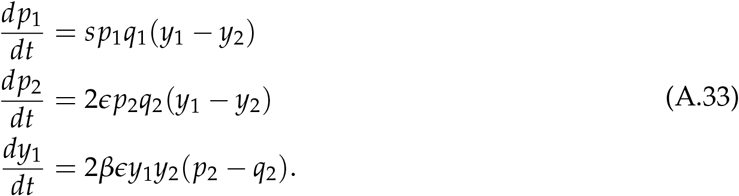

This model is the same as A.19. This explains why this variation of the full model has dynamics like that of the original H-GM model.

##### H-GM Plant Resource Dependent Model without Re-assortment

In the above models, the qualitative dynamics are exactly the same, even down to the eigenvalues. Considered next is the model variation in which mating and egg-laying were done on the same plant without the re-assortment of females. Qualitative dynamics appear to be the same from numerical simulations, but the analytical advantages are not significant. Therefore, this model is less useful because this simplification requires a biologically less realistic assumption.

Using approximation techniques to the H-GM plant resource dependent model without reassortment, the following equations are derived:

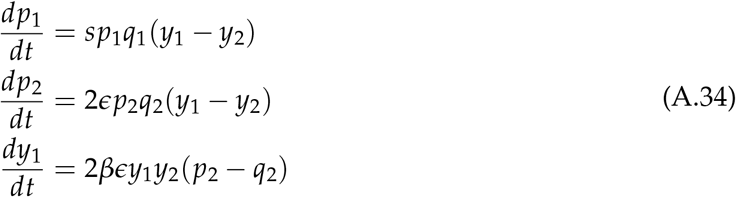

This model again reduces to A.19. This explains why this variation of the full model also has dynamics like that of the original H-GM model.

## Appendix B: Fig-fig Wasp Model

### Dynamics of Insects

In *Ficus-Courtella wardi* (Wade, 2007), the female enters a female phase fig, but in doing so loses her wings and dies within the fig after laying eggs in each ovule. During the interfloral phase of the fig, the fig wasp larvae mature then mate within the fig. When the fig reaches the male phase, the male fig wasps chew an opening for the females, and the females leave the fig carrying pollen. The plant genetics, however, become much more complicated. The male phase of the fig from which females emerge, laden with eggs, must overlap with the female phase (the start of a new floral generation of another fig) so that the fig wasp can pollinate the fig and oviposit her eggs. These overlapping generations are a major biological difference from the H-GM system.

The frequency of adult insects of type *i* found on plants of type *m* is denoted by *x*_*i,m*_ ^1^. Let *y*_*m*_ and *z*_*m*_ be the frequencies of the types of plants during the insect mating period in male and female phase during even during odd generations, respectively, and female and male phase during even generations, respectively. Again, indices *i, j, k* are used for insects and *l, m, n* for plants. Assuming random mating on host, the frequency of mating pairs formed by females *i* and males *j* on a plant of type *m* is

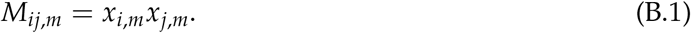

Therefore, the frequency of eggs with genotype *k* produced by pairs (*i, j*) mating on a plant of type *m* is *M*_*ij,m*_*R*(*i, j* → *k*), where *R* gives the corresponding recombination frequencies for the genotypes in this system.

Females leave the fig carrying eggs^2^. Assume each fig has a carrying capacity of wasps that it can support. Therefore, the frequency of females leaving a particular fig, *i*.*e*. the contribution of each mating pool to offspring, is equal to the proportion of the fig type in that system. This is a reasonable assumption since there is a limited resource inside the fig which leads to larval competitionWeiblen (2002). Females leave the male phase figs and bring pollen to female phase figs. The proportion of eggs with genotype *k* and carried by mothers with genotype *i* in the whole set of eggs before oviposition is

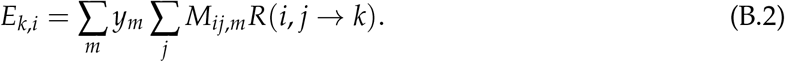

Females then search for a suitable female phase fig in which to oviposit the eggs. The frequency of eggs with which genotype *k* oviposited on a plant of type *n* is

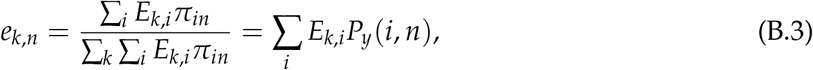

where

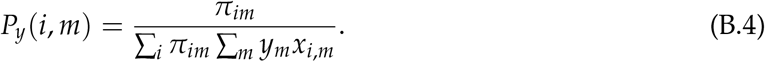

After selection on a plant of type *n*, the frequencies of larvae in the plant are

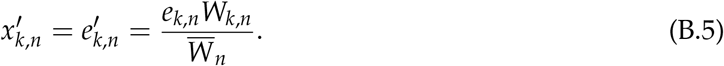

Recall it is assumed that the percentage contribution of each larva pool (*i*.*e*. plant) to the total larvae population is equal to the proportion of each plant type. Therefore the final equation for insects going from the *Y* population of plants to the *Z* population of plants is

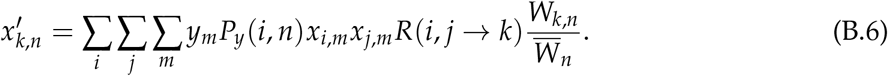

Two plant populations need to be fertilized consecutively to complete a full plant generation of both male and female phase plant ^3^. Therefore, two generations of insects are modeled consecutively. The second generation is calculated in the same way above except the plant population in the male phase is *Z*, so where *y* was used, now *z* is used.

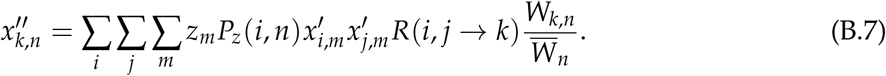

The probability that a female of type *i* goes to a plant of type *n* to lay the eggs is

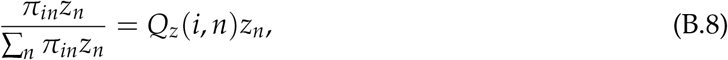

where

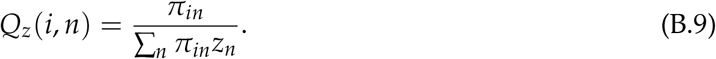

The female insect pollinates a plant *n* in population *Z* by pollen from plant *m* in population *y* with probability

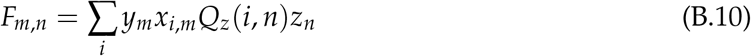

which can be interpreted as mating probability of male plant type *m* from the *Y* population of fig trees and female plant type *n* from the *z* population of fig trees. Thus, the frequency of plant *l* produced as a result of mating of plants *m* and *n* is *F*_*mn*_*S*(*m, n* → *l*), where *S*(*m, n* → *l*) is the corresponding segregation probability.

Assume flowering time is paternally inherited, *i*.*e*. the offspring of these matings from the *Y* population to the *Z* population will contribute offspring only to the *Y* flowering population. So the frequency of offspring is

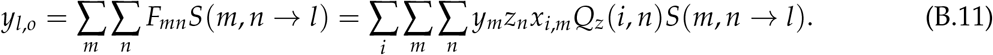

Finally, to account for the fact that figs last more than one generation, let only a proportion *β* of plants be replaced each generation. Then

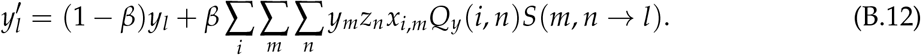

The model developed thus far models *Y* pollinating *Z*. Modeling the next generation of fig wasp pollination, when *Z* pollinating *Y*, leads to a final equation of

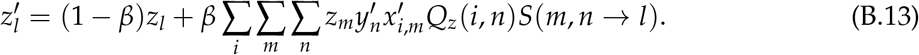

## Footnotes

1 Note: This model starts with the part of the insect life-cycle after larval selection. So if we assume all loci under consideration are diallelic, then we have 8 insect types to consider. We also now have 4 types of plants, 2 types in each population. This contrasts the H-GM model in which we started the life cycle with the females in the air assorting to plants to mate. In the fig wasp, most of the life cycle takes place within one fig (so 8 cases between the two fig types currently harboring a population), and when the female leaves, she leaves with eggs (so 16 cases).

2 This is the only time females are in the air, leaving the host plant. This is unlike the original Greya moths which required they mate and lay eggs on different plants, assorting multiple times.

3 3To model the second plant population, one more insect generation needs to be modeled. This second insect generation will depend on *z*, as Z is now the plant population that is in the male phase.

